# AAV-mediated gene transfer restores a normal muscle transcriptome in a canine model of X-linked myotubular myopathy

**DOI:** 10.1101/499384

**Authors:** Jean-Baptiste Dupont, Jianjun Guo, Michael W. Lawlor, Robert W. Grange, John T. Gray, Ana Buj-Bello, Martin K. Childers, David L. Mack

## Abstract

Multiple clinical trials employing recombinant adeno-associated viral (rAAV) vectors have been initiated for neuromuscular disorders, including Duchenne and limb-girdle muscular dystrophies, spinal muscular atrophy, and recently X-linked myotubular myopathy (XLMTM). Previous work from our laboratory on a canine model of XLMTM showed that a single rAAV8-cMTM1 systemic infusion corrects structural abnormalities within the muscle and restores contractile function, with affected dogs surviving more than four years post injection. This exceptional therapeutic efficacy presents a unique opportunity to identify the downstream molecular drivers of XLMTM pathology, and to what extent the whole muscle transcriptome is restored to normal after gene transfer. Herein, RNA-sequencing was used to examine the transcriptomes of the *Biceps femoris* and *Vastus lateralis* in a previously-described canine cohort showing dose-dependent clinical improvements after rAAV8-cMTM1 gene transfer. Our analysis confirmed several dysregulated genes previously observed in XLMTM mice, but also identified new transcripts linked to XLMTM pathology. We demonstrated XLMTM transcriptome remodeling and dose-dependent normalization of gene expression after gene transfer and created new metrics to pinpoint potential biomarkers of disease progression and correction.

## Introduction

The last 30 years have seen the field of neuromuscular disease (NMD) genetics grow from one known gene, for Duchenne muscular dystrophy (DMD) in 1987, to hundreds of disease genes identified today. Almost all NMDs are classified as rare conditions, but collectively they are not rare, creating a serious public health problem and an enormous burden on the patients and families affected by these debilitating disorders. The standard of care for NMDs consists almost entirely of rehabilitation modalities to help patients cope with their deteriorating condition, surgical procedures, and steroids in specific cases (Bushby, Finkel et al., 2010a, Bushby, Finkel et al., 2010b, Wang, Bonnemann et al., 2010, Wang, Dowling et al., 2012, Wang, Finkel et al., 2007). However, the future for a subset of these NMD patients looks brighter, as several phase I/II gene therapy clinical trials have reported remarkable safety and efficacy outcome measures. NMDs are complex diseases whose pathological processes are often not entirely understood. Gene therapy offers a unique treatment approach for single-gene NMDs, because fixing the root cause of the defect does not necessarily require a comprehensive understanding of the pathophysiology. However, successful gene therapy provides an extraordinary opportunity to delve more deeply into the complex molecular underpinnings of disease initiation, progression and correction.

In the past years, treatment breakthroughs have been made in small and large animal models of NMDs (Bengtsson, Hall et al., 2017a, Bengtsson, Hall et al., 2017b, Childers, Joubert et al., 2014, Le Guiner, Servais et al., 2017, Mack, Poulard et al., 2017, Malerba, Klein et al., 2017, Pozsgai, Griffin et al., 2017) and clinical benefits have been reported after a single injection of recombinant adeno-associated virus (rAAV) in patients with spinal muscular atrophy (Mendell, Al-Zaidy et al., 2017). In most of these studies, results consist of correlations between transgene expression levels and clinical outcome measures. However, this level of insight may prove insufficient when sub-therapeutic effects are the outcome, or conversely when adverse events occur *in vivo* (Gicquel, Maizonnier et al., 2017, Mendell, Campbell et al., 2010, Thomsen, Alkaslasi et al., 2017), strongly arguing for a deeper understanding of disease mechanisms. Therefore, the identification of disease-associated gene dysregulation and signalling pathways that are not – or only partially – corrected after gene transfer is of considerable interest. Also, as more gene therapies are approved for NMDs, it will become increasingly important to track their long-term durability using non-invasive biomarkers that appear prior to clinical deterioration.

Recent advances in -omics technologies have provided valuable tools for the diagnosis of NMDs and for the understanding of disease mechanisms at the molecular level (Butterfield, Dunn et al., 2017, Cummings, Marshall et al., 2017, Gama-Carvalho, M et al., 2017, Liang, Tian et al., 2017a, Liang, Tian et al., 2017b). In this study, we investigated the underlying mechanisms of rAAV-mediated correction of muscle pathology utilizing the canine model of X-linked myotubular myopathy. XLMTM is a centronuclear myopathy caused by mutations in the *MTM1* gene, coding for the myotubularin protein (Laporte, Hu et al., 1996). Patients exhibit severe muscle weakness, breathing difficulties, and a median survival of 1-2 years (Beggs, Byrne et al., 2018). XLMTM muscle cells have smaller diameters, an abnormal architecture, and a poor excitation-contraction coupling (ECC) capacity (Al-Qusairi, Weiss et al., 2009, Kutchukian, Lo Scrudato et al., 2016, Lawlor, Beggs et al., 2016, Shichiji, Biancalana et al., 2013). During the past decade, our lab and others have used rAAV vectors in XLMTM mice and dogs and demonstrated remarkable therapeutic efficacy, paving the way for a recently initiated clinical trial (Buj-Bello, Fougerousse et al., 2008, Buj-Bello, Laugel et al., 2002, Childers et al., 2014, Mack et al., 2017), (ClinicalTrials.gov identifier NCT03199469). Treated XLMTM dogs showed dose-dependent correction of multiple disease phenotypes, resulting in improved muscle function and survival up to 4 years of age (Elverman, Goddard et al., 2017, Mack et al., 2017). Here, we explored the transcriptional changes driving this correction of muscle morphology and function in treated XLMTM dogs using an unbiased and comprehensive approach based on RNA-sequencing (RNA-seq). Untreated XLMTM dogs exhibited transcriptional changes consistent with well-described disease phenotypes, and several genes emerged as new potential candidates to explain disease development and progression. Importantly, rAAV gene therapy lead to a dose-dependent restoration of the muscle transcriptome toward a more normal profile. We also present an array of metrics based on RNA-seq data to quantify gene therapy efficacy and distinguish the transcripts corrected to normal after treatment from those escaping correction. Finally, we highlight potential RNA biomarkers to monitor and predict the efficacy of current and future gene therapy protocols.

## Results

XLMTM dogs were previously randomised to receive any of the three increasing doses of rAAV8-*cMTM1*, or saline as a negative control. In this study, these groups were named **AAVLow** (0.3E+14 vector genomes per kilogram, vg/kg), **AAVMid** (2.0E+14 vg/kg), **AAVHigh** (5.0E+14 vg/kg), and **XLMTM**, respectively, and were compared to wild-type (**Healthy**) controls (Fig 1A). In a previous study, the AAVMid and AAVHigh doses were characterized as therapeutic with full correction of disease phenotypes such as survival, neurological scores, muscular and respiratory functions (Fig 1B, (Mack et al., 2017). Conversely, the AAVLow dose resulted in sub-therapeutic clinical outcomes, including significant weakness, significant muscle pathology, and death of two animals. In this *a posteriori* study, the transcriptome of these dogs was analysed using RNA-seq on two distinct muscles: *Biceps femoris*, and *Vastus lateralis*. Due to premature death encountered in the XLMTM and AAVLow groups, dogs were not age-matched, but samples were all collected at necropsy to reflect the final extent of disease progression or correction (**Appendix Table 1**). In each muscle, the top 500 most informative genes were selected based on their expression variance across samples and were used for principal component analysis (PCA). It showed segregation of the 13 dogs in 2 groups (Fig 1C-D): AAVLow and XLMTM dogs in one group; Healthy and AAVHigh dogs in the other. The AAVMid group also clustered with Healthy and AAVHigh dogs in the *Vastus*, and was spread between the two groups in the *Biceps*. Gene ontology analysis was performed on the top 500 genes in each muscle, and identified multiple terms related to muscle biology, particularly in the *Vastus* (**Appendix Figure 1**).

**Figure 1:**
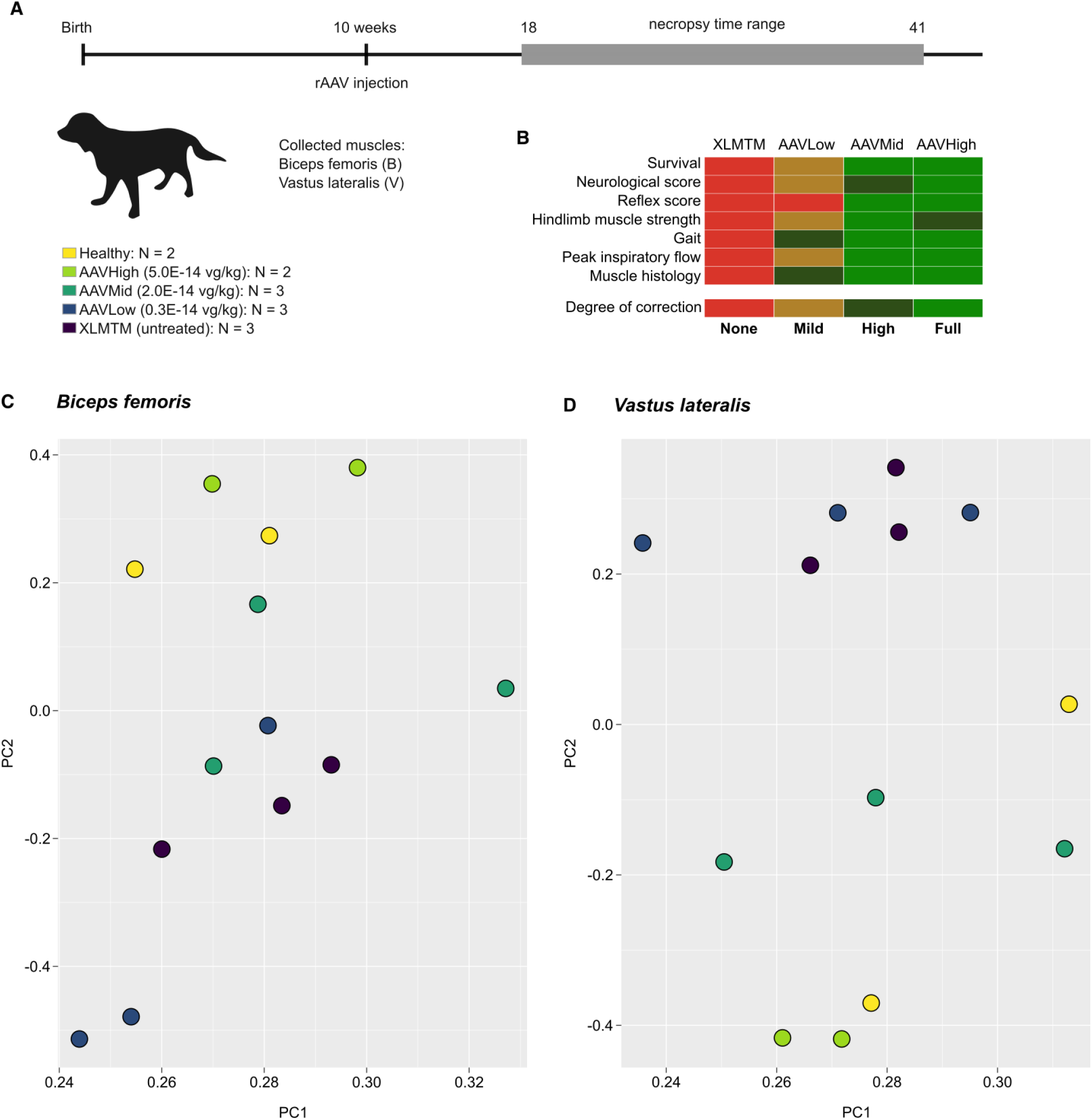
Visualisation of gene therapy efficacy by transcriptome profiling in XLMTM dogs. (**A**) Experimental timeline in weeks and details of the 5 groups of dogs. The rAAV vector was injected at 10 weeks of age, and dogs were monitored until the age of 39 to 41 weeks, unless euthanasia criteria were reached prematurely, as encountered in untreated XLMTM and AAVLo dogs. Additional details are provided in Appendix Table 1. (**B**) Heatmap representing the degree of corrections of key XLMTM phenotypes achieved in the three groups of dogs, as measured in our previous study (Mack et al., 2017). (**C**) Principal Component Analysis (PCA) on RNA-seq data in the *Biceps femoris* after selection of the top 500 most informative genes across samples. (**D**) Principal Component Analysis on RNA-seq data in the *Vastus lateralis* after selection of the top 500 most informative genes across samples. For **C** and **D**, PC1 and PC2 were projected in a twodimensional plot, with each dog represented as a color-coded dot.

To gain further insight into the development of XLMTM disease phenotypes in these two muscles, and their correction by gene therapy, the differentially expressed (DE) genes between XLMTM dogs and healthy controls were determined using the edgeR package (Robinson, McCarthy et al., 2010). In the *Biceps* and the *Vastus*, 824 and 1122 DE genes were obtained, respectively. Among them, 400 were identified in both muscles, suggesting overlap in XLMTM disease process between different muscles. Gene ontology analysis of the 400 common DE genes pointed to muscle development, muscle contraction, and various metabolic responses (Fig 2A). The lists of DE genes in both *Biceps* and *Vastus* are available in **Appendix Tables 2 and 3**. To visualize the extent of XLMTM transcriptome remodelling achieved by the three injected rAAV doses, the entire set of DE transcripts from both muscles of XLMTM untreated dogs and all three treatment groups were plotted relative to healthy control dogs (log fold-change values, log FC) (Fig 2B-C). This plot showed that both up-regulated and down-regulated DE genes (in green and blue, respectively) were corrected, with log FC values progressively returning to zero. The log FC interquartile (IQ) range was then calculated to quantify transcriptome remodelling. This number represents the length of the interval comprising the central half (25%-75%) of ranked log FC values. Therefore, it estimates how the expression values of the DE gene set diverge from healthy controls. In the *Biceps*, log FC IQ range values varied from 3.0 to 2.8, 1.3, and 0.91 in untreated XLMTM, AAVLow, AAVMid and AAVHigh dogs, respectively (Fig 2B). In the *Vastus*, AAVMid dogs had a lower IQ range than AAVHigh dogs, reflecting a higher transcriptome remodelling effect with the intermediate rAAV dose (XLMTM: 2.7, AAVLow: 2.1, AAVMid: 0.81, and AAVHigh: 1.3, Fig 2C). Hierarchical clustering using the common DE genes was able to distinguish therapeutic (AAVMid and AAVHigh) and sub-therapeutic (AAVLow) doses (Fig 2D). The only exception was observed between Dog 04 (AAVMid) and Dog 13 (AAVLow) *Biceps* samples, which clustered with the sub-therapeutic and therapeutic groups, respectively. Interestingly, disease correction in Dog 13 (SSAN_18) was more pronounced than in the two other AAVLow dogs, and it did not reach euthanasia criteria before the end of the study period (**Appendix Table 1**, (Mack et al., 2017). *MTM1* transgene expression was then measured directly from RNA-seq data using a custom-made workflow for rAAV-derived mRNA tag counting and showed that transgene expression strongly correlated with the extent of transcriptome correction (Fig 2D). Transgene expression level in Dog 13 *Biceps* was comparable to that of AAVMid dogs (*MTM1* tag count = 1.3 vs. a median of 1.2 in AAVMid dogs), suggesting that evaluation of transgene expression directly from RNA-seq data is a better predictor of transcriptome remodelling than the injected dose of gene therapy vector, and that it reflects itself on overall treatment outcome. Therefore, injected *Biceps* samples were re-ordered in two distinct groups: the *Therapeutic* group with the two AAVHigh dogs, two AAVMid dogs, and Dog 13; and the *Sub-therapeutic* group, with the two remaining AAVLow dogs, and Dog 04. In the *Vastus*, all AAVMid and AAVHigh clustered together with Healthy controls, and formed the *Therapeutic* group, while AAVLow dogs formed the *Sub-therapeutic* group.

**Figure 2:**
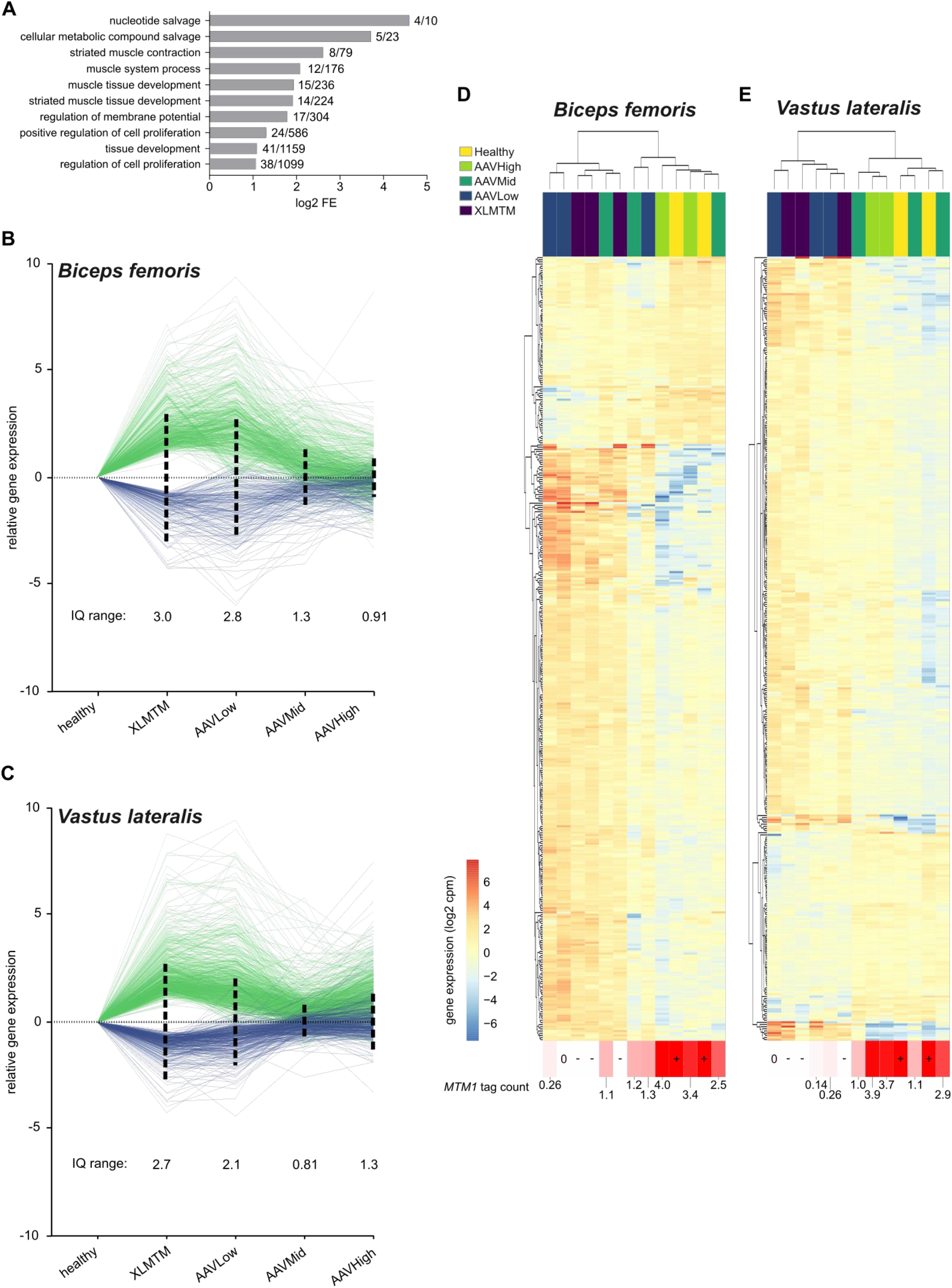
Remodeling of the XLMTM transcriptome by rAAV gene therapy. (**A**) Gene ontology analysis on the differentially expressed genes common to XLMTM *Biceps* and *Vastus* muscles. For each represented term, the log2 fold enrichment (FE) values are graphically represented, together with the x/y ratio, defined as the number of genes participating to this enrichment (x) over the total number of genes affiliated with the corresponding ontology (y). (**B**) Relative expression of the differentially expressed genes in the *Biceps*, expressed in log fold change (log FC) after normalization with healthy controls. The interquartile ranges were determined from positive log FC values (negative log FC were converted into absolute values) and represented as dashed vertical lines pointing in both directions around the y=0 line. (**C**) Similar representation for the differentially expressed genes detected in the *Vastus* muscle. (**D-E**) Hierarchical clustering generated from log cpm values of the 400 common differentially expressed genes for the *Biceps* (**D**) and the *Vastus* (**E**). Each column represents one muscle sample, color-coded for treatment group; each row represents one individual gene, color coded for its expression level (log cpm). RNA-seq-based *MTM1* mRNA tag counts are indicted under each column.

Since distinct sub-populations of transcripts may respond differently to gene therapy, an array of new metrics was created, based on the type of transcriptome remodelling effect. *Rescued* genes were defined as the XLMTM-associated DE genes that were no longer significantly different from healthy controls in rAAV-treated dogs (Fig 3A). Conversely, *Resistant* genes are still differentially expressed between rAAV-treated dogs and healthy controls. Two additional subpopulations of *Partially Rescued* and *Worsened* genes were also defined but were both marginally represented irrespective of the vector dose. These metrics were determined for the *Sub-therapeutic* and *Therapeutic* groups of dogs, in the *Biceps* and the *Vastus* muscles (Fig 3B-C). Remarkably, a sub-therapeutic effect was associated with only 6.1 % and 7.5 % of genes with a *Rescued* expression profile in the *Biceps* and *Vastus* muscles, respectively, while *Resistant* genes accounted for 64 % and 57 %. In sharp contrast, the proportion of *Rescued* genes reached 52.1 % and 42.7 % in the *Biceps* and *Vastus* of the dogs showing therapeutic benefits. This indicates that an increase in *MTM1* transgene expression drives a more profound restoration of the muscle transcriptome, providing for the first time a comprehensive transcriptomic explanation of physiological outcome measures observed previously (Mack et al., 2017). Next, populations of *Candidate Biomarkers* of XLMTM correction were defined as the genes presenting a *Rescued* expression profile in the *Therapeutic*, but not in the *Sub-therapeutic* group of dogs (Fig 3B-C, right panel). This resulted in more than 400 genes in each muscle, 120 of which were detected in both *Biceps* and *Vastus*. As expected, the expression pattern of these 120 genes followed a similar trend, with sustained deregulation in the XLMTM and *Sub-therapeutic* groups, and a marked rescue in the *Therapeutic* group (**Appendix Figure 3**). In the latter, IQR values were only 0.45 and 0.61 in the *Biceps* and the *Vastus*, respectively. Finally, a list of twelve candidate biomarkers was isolated among the genes with the most significant expression changes in the two XLMTM muscles (P-value < 0.001 in each muscle, Fig 3D) and their expression level was correlated with *MTM1* tag counts in injected dogs and untreated controls (tag counts = 0, by definition). The most significant correlation was obtained for *APEX2*, an endonuclease involved in base-excision-repair and progression through the cell cycle (ρ = -0.86, P-value = 2.3E-7). Interestingly, this list also contained *ALAS2* (r = -0.83, P-value = 1.3E-6) and *PIK3R2* (r = -0.82, P-value = 3.3E-6), two genes that were recently associated with muscle strength (Pilling, Joehanes et al., 2016), and *NRK* (r = -0.82, P-value = 3.8E-6), known to be specifically expressed during myotome formation and early muscle development in mice, but not in the adult (Kanai-Azuma, Kanai et al., 1999). The expression of these 4 genes in injected dogs was inversely correlated with *MTM1* transgene expression. *Candidate Biomarkers* also included *RIN1, EFCAB7, ANGPTL2, CHRNA1, PROCR, IGF2, ETS2*, and *RIMS3* (Fig 3E). The full lists of *Rescued* genes in both muscles and of *Candidate Biomarkers* are available in **Appendix Table 4 and Table 5**.

**Figure 3:**
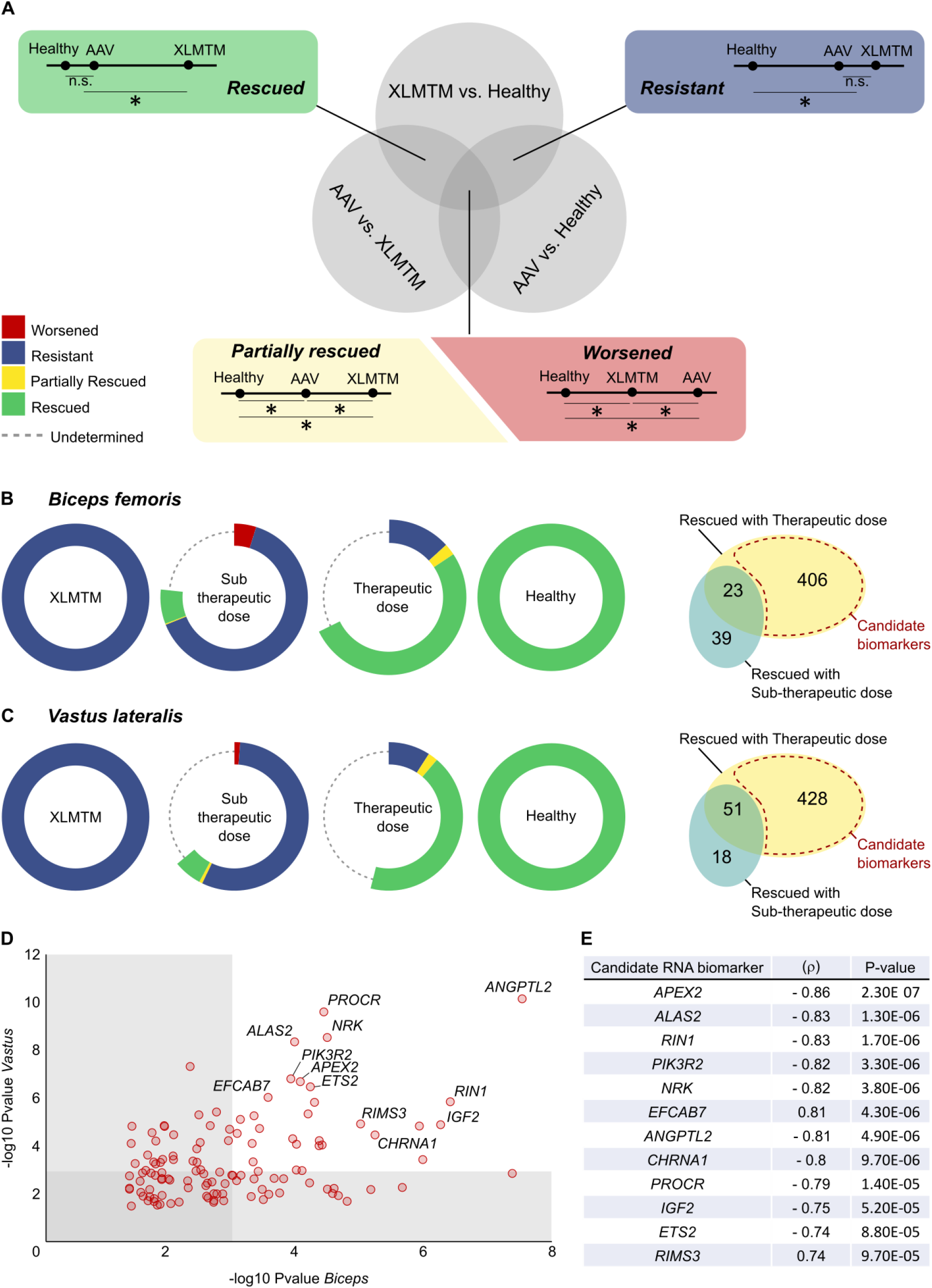
Transcriptome remodelling metrics and candidate RNA biomarkers of gene therapy efficacy in XLMTM dogs. (**A**) Venn diagram representing the intersections of differentially expressed gene populations obtained after comparison of untreated XLMTM, rAAV-treated XLMTM, and healthy dogs. Focus is made on the *Rescued* and the *Resistant* two-circle domains in specific insets, and on the central domain, containing *Partially Rescued* and *Worsened* genes. Exemplar gene expression levels are plotted on a line for the three groups: healthy, XLMTM and rAAV-treated (AAV) dogs. N.s. non-significant gene expression difference, * significant difference. (**B-C**) Evolution of the described metrics in the *Sub-therapeutic* and the *Therapeutic* groups of dogs – as defined in the text – in the *Biceps* (**B**) and the *Vastus* (**C**) muscles. The initial DE gene set in XLMTM dogs is represented as a blue ring, and its counterpart in healthy controls as a green ring. In rAAV-treated dogs, the areas corresponding to the above-described metrics have an arc length directly proportional to the proportion of initial DE genes presenting the corresponding expression profile. The right panels represent the Venn diagrams showing the number of genes with a *Rescued* profile and their overlap between the *Sub-therapeutic* and the *Therapeutic* groups. The *Candidate Biomarker* gene subsets are delineated with a dashed red area. (**D**) Two-dimensional dot-plot of the 120 candidate biomarkers common to the *Biceps* and *Vastus* muscles, representing the significance of the differential expression between XLMTM dogs and Healthy controls (P-values). Each dot represents one gene. (**E**) Selection of twelve RNA biomarkers of gene therapy efficacy in XLMTM dogs, ranked according to their correlation coefficient (ρ) when compared to *MTM1* mRNA tag counts.

## Discussion

With the recent success of preclinical and clinical trials using rAAV vectors, understanding the exact processes by which disease phenotypes are impacted – and eventually rescued – by gene therapy is of considerable interest for the field of molecular medicine. Here, we take advantage of the unprecedented therapeutic efficacy obtained in XLMTM dogs to investigate the molecular mechanisms of NMD correction, with a focus on pathological muscle transcriptome. Overall, our study demonstrates that corrective rAAV gene therapy induces a dose-dependent transcriptome remodelling, which was highly correlated with RNA-seq-based transgene expression measurements. These measures were comparable with our previous study despite being obtained by different techniques on distinct muscle biopsies (Mack et al., 2017) and revealed high levels of recombinant *MTM1* transcript in the dogs presenting with full disease correction.

To our knowledge, this is the first reported comprehensive transcriptome profiling from myotubularin-deficient muscles. Previously, individual genes were queried in *Mtm1*-deficient mice as an attempt to explain clinical observations, (Al-Qusairi, Prokic et al., 2013, Dowling, Joubert et al., 2012, Mariot, Joubert et al., 2017). In addition, muscle biopsies from 8 XLMTM patients have been analysed for the expression of 4,200 genes using microarray and pointed to the upregulation of genes coding for cytoskeletal and extracellular matrix proteins (Noguchi, Fujita et al., 2005). In our study, the use of RNA-seq allowed us to gather data on more than 32,000 dog genes, some of them not yet characterized. Importantly, several genes previously identified as differentially expressed in XLMTM patients and XLMTM mice were also detected in dogs. This notably includes three subunits of the nicotinic acetylcholine receptor: *CHRNA1*, which is up-regulated in the three species, *CHRNG*, also up-regulated in XLMTM dogs and mice, and *CHRND*, up-regulated in dogs and mice but down-regulated in XLMTM patients. This deregulation of nicotinic acetylcholine receptor genes directly reflects the inefficacy of XLMTM muscle cells to generate Ca2+ transients in response to electrical stimulation (Al-Qusairi et al., 2009, Kutchukian et al., 2016). Interestingly, XLMTM dogs also showed deregulated expression of multiple additional genes coding for ion transporters, that could also lead to ECC defects, including sodium (*SCN4B, SCN5A, SCNN1A*) and potassium (*KCNK17, KCNQ5*) channels, the sodium/potassium-transporting ATPase subunit alpha-3 (*ATP1A3*), and multiple members of the solute carrier (SLC) family. Another major consequences of myotubularin deficiency is an early muscle hypotrophy observed in patients and animal models (Beggs, Bohm et al., 2010, Buj-Bello et al., 2002, Goddard, Mack et al., 2015, Lawlor et al., 2016, Pierson, Dulin-Smith et al., 2012, Shichiji et al., 2013). XLMTM myotubes have a small diameter and centrally positioned nuclei, and they are often considered abnormally developed or unable to maintain a fully differentiated state. In XLMTM mice, this has been correlated with deregulated PI3K/Akt/mTOR signalling and abnormal autophagy (Al-Qusairi et al., 2013, Fetalvero, Yu et al., 2013, Joubert, Vignaud et al., 2013, Lawlor, Viola et al., 2014). More recently, another study reported the downregulation of the myostatin pathway in *Mtm1*-KO mice, which could be a compensatory mechanism to counter muscle hypotrophy (Mariot et al., 2017). Here, the genes coding for myostatin (*MSTN*) and follistatin (*FST*) were respectively found down– and up-regulated in XLMTM dogs, confirming the involvement of this pathway downstream of myotubularin deficiency (**Appendix Figure 2**), and its potential as a therapeutic target. Indeed, treatment with ActRIIB-mFc, a myostatin inhibitor, resulted in muscle hypertrophy and increased lifespan in XLMTM mice (Lawlor, Read et al., 2011). More recently, these mice were co-injected with a rAAV vector coding for an inhibitory myostatin propeptide in addition to rAAV-*Mtm1*, and this led to a further increase in muscle mass than mice treated with rAAV-*Mtm1* only (Mariot et al., 2017). This demonstrates that the tools created in this study have the potential to identify relevant targets for combinatorial gene therapy applications. In the light of our RNA-seq data, abnormal muscle development appears as an important component of XLMTM biology in dogs. The genes encoding the myogenic regulators MyoD (*MYOD1*) and myogenin (*MYOG*) are over-expressed in XLMTM dogs, together with *PAX7*, a marker of adult muscle stem (satellite) cells (**Appendix Tables 2 & 3**). In addition, several genes coding for isoforms of structural proteins transiently expressed during skeletal muscle embryonic development were found up-regulated, including the acetylcholine receptor γ subunit (*CHRNG*) and myosin heavy chain 3 (*MYH3*) in the *Biceps*, or cardiac alpha-actin (*ACTC1*) and troponin T (*TNNT2*) in the *Vastus*. Overall, transcriptome profiling of canine XLMTM muscles has not only confirmed gene expression data in mice, but also helped identify multiple genes never associated with XLMTM biology, whose deregulation gives a new insight into disease phenotypes.

Gene therapy studies rarely investigate the molecular consequences of target cell transduction. With rAAV vectors injected in new patients every year, now including boys suffering from XLMTM (clinical trial NCT03199469), a better understanding of the molecular mechanisms by which NMDs are corrected will become crucial. Recently, one study reported the normalization of microarray gene expression data after dual rAAV gene therapy in a mouse model of oculopharyngeal muscular dystrophy (OPMD), (Malerba et al., 2017), but the authors did not focus on specific genes or gene sub-populations. Here, we provide for the first time a comprehensive picture of muscle transcriptome remodelling after rAAV gene therapy using RNA-sequencing. We used hierarchical clustering to group injected dogs based on the extent of transcriptome remodelling rather than on the dose of rAAV injected *in vivo*. In agreement with previous clinical observations (Mack et al., 2017), this resulted in a *Sub-therapeutic* and a *Therapeutic* group of dogs. Strikingly, the *Biceps* sample collected from Dog 13 was part of the *Therapeutic* group despite a treatment with an otherwise sub-therapeutic dose of vector. However, this correlates with the extended lifespan and phenotypic improvements observed in this specific dog (Mack et al., 2017). To illustrate different facets of transcriptome remodelling, we created several metrics and determined the proportion of genes whose expression was normalized or not after treatment (Fig 3A). We demonstrated that these metrics can be used to identify dogs exhibiting therapeutic benefits from those in which gene therapy efficacy was not durable.

In addition, this approach proved very effective to identify RNA biomarkers of XLMTM correction. Interestingly, several of the top twelve candidate genes are not only in line with XLMTM phenotypes but could also help deepen our understanding of disease development: *ALAS2* and *PIK3R2* have recently been negatively associated with muscle strength in a meta-analysis in more than 7,500 human subjects (Pilling et al., 2016), and were also found over-expressed in XLMTM dogs; Nik-related kinase (NRK) is a protein kinase involved in actin polymerisation through phosphorylation and inhibition of cofilin is expressed only during embryonic myogenesis (Kanai-Azuma et al., 1999), and *NRK* is overexpressed in XLMTM dogs, which adds on the list of deregulated developmental genes; *APEX2* encodes an endonuclease physically associated with proliferating cell nuclear antigen (PCNA) and driving cell cycle progression through base-excision-repair on the replicative nuclear and mitochondrial DNA (Ide, Tsuchimoto et al., 2004, Tsuchimoto, Sakai et al., 2001). *APEX2* is also over-expressed in untreated XLMTM muscles and corrected in the *Therapeutic* group, suggesting that de-regulation / re-regulation of the cell cycle is a key component of disease development and rescue. Altogether our data provide a transcriptional characterization of the structural immaturity encountered in XLMTM muscles. RNA biomarkers defined in this study could also help identify protein products known to be secreted in the blood, and therefore, easily quantifiable. In this regard, at least two genes highlighted in our list could be further investigated at the protein level: angiopoietin-like protein 2 (*ANGPTL2*), an inflammatory mediator participating in blood vessel formation, and insulin-like growth factor 2 (*IGF2*). Recently, increased *ANGPTL2* expression has been reported in a mouse model of denervation-induced muscle atrophy, and its knockout was able to improve muscle growth and satellite cell activity (Zhao, Tian et al., 2018). Here, *ANGPTL2* is over-expressed in untreated XLMTM dogs, and reduced to basal level in the *Therapeutic* group. Thus, ANGPTL2 itself might be an important mediator of rAAV-mediated therapeutic benefits. As a proinflammatory cytokine, it is accurately quantifiable in blood samples, and is already used in the context of colorectal cancer (Toiyama, Tanaka et al., 2014). This makes it an excellent candidate biomarker to monitor the evolution of XLMTM after clinical gene therapy.

In conclusion, this study brings evidence of XLMTM rescue by rAAV gene therapy at the molecular level, in the form of a dose-dependent transcriptome remodelling acting on specific aspects of disease biology. Furthermore, it introduces an array of RNA biomarkers in the form of individual genes or carefully defined gene populations, which brings unprecedented insight to the mechanisms of NMD rescue by gene therapy.

## Supporting information

## Materials and Methods

### Animal model

The XLMTM Labrador/Beagles used in this study bear a p.N155K mutation in the *MTM1* gene, and were issued from a colony maintained at the University of Washington (Beggs et al., 2010, Childers et al., 2014). More precisely, they were previously included in a gene therapy study, the methods and results of which were extensively described in a recent publication (Mack et al., 2017). In brief, XLMTM dogs were infused with three increasing doses of a rAAV vector expressing a canine MTM1 cDNA: 0.3E+14, 2.0E+14, and 5.0E+14 vg/kg at the age of 10 weeks, and sacrificed between 39 and 41 weeks old, or when reaching humane euthanasia criteria. In the absence of treatment, XLMTM dogs, both hemizygous males and homozygous females, exhibit progressive clinical deterioration between about 12 and 26 weeks of age, resulting in an inability to stand, walk, and feed. Eight injected dogs were included in this study, together with 3 untreated XLMTM and 2 Healthy controls, as summarized in **Appendix Table 1**.

### RNA extraction and sequencing

Muscle necropsy tissues were collected, frozen on dry ice, and stored at ≤-60°C. RNA extraction, library preparation and sequencing on Illumina Hiseq 4000 were conducted by Genewiz Inc (NJ, USA). Specifically, total RNA was extracted with Qiagen AllPrep mini kit. RNA samples were quantified using Qubit 2.0 Fluorometer (Life Technologies, Carlsbad, CA, USA) and RNA integrity was checked with RNA Screen Tape on Agilent 2200 TapeStation (Agilent Technologies, Palo Alto, CA, USA). RNA sequencing library preparation was prepared with TruSeq Stranded mRNA library Prep kit following manufacturer’s protocol (Illumina, Cat# RS-122-2101). Sequencing libraries were validated using DNA Analysis Screen Tape on the Agilent 2200 TapeStation (Agilent Technologies, Palo Alto, CA, USA), and quantified by using Qubit 2.0 Fluorometer (Invitrogen, Carlsbad, CA) as well as by quantitative PCR (Applied Biosystems, Carlsbad, CA, USA). Sequencing libraries were multiplexed and clustered on flow cell using the cBOT from Illumina. After clustering, flow cell was loaded on the Illumina HiSeq instrument according to manufacturer’s instructions. The samples were sequenced using a 2×150 Pair-End (PE) High Output configuration. Image analysis and base calling were conducted by the HiSeq Control Software (HCS) on the HiSeq instrument. Raw sequence data (.bcl files) generated from Illumina HiSeq was converted into fastq files and demultiplexed using Illumina bcl2fastq program version 2.17. One mismatch was allowed for index sequence identification. The 13 *Vastus*-mRNA-derived libraries were sequenced to an average of 60M paired-end reads/sample and the 13 *Biceps*-mRNA-derived libraries were sequenced to an average of 20M paired-end reads/sample.

### Data analysis

Initial data analysis, including alignment to reference genome and count table generation, was conducted on the DNAnexus platform (Palo Alto, CA, USA). Specifically, fastq files were first mapped onto the reference genome of *Canis lupus familiaris* (Genome assembly: CanFam3.1 (GCA_000002285.2), downloaded from Ensembl.org) using the Subread mapping package (Liao, Smyth et al., 2013, Liao, Smyth et al., 2014) (settings: subread-align -t 0 -T 4 -d 50 -D 600 -i my_index -r read1 -S ff -R read2 -o $output_name). The resulting bam files were fed into featureCounts (Liao et al., 2014) package to obtain the count-per-gene table (settings: featureCounts -p -P -d 50 -D 600 -t exon -g gene_id -a GTF_file -o $output_name bam_file1 bam_file2 bam_file3 bam_file4 bam_file5 bam_file6 bam_file7 bam_file8 bam_file9 bam_file10 bam_file11 bam_file12 bam_file13). Count tables were further analysed with the open-source RStudio environment for R (https://www.r-project.org/), and the Bioconductor software https://www.bioconductor.org/. The limma (Ritchie, Phipson et al., 2015) and edgeR (Robinson et al., 2010) packages were used to normalize, fit, and compare the data between groups following the analysis pipeline detailed here (Law, Alhamdoosh et al., 2016). Cutoff values for DE gene determination were as follows: Pvalue < 0.05, and fold-change (FC) > 2.0. The code used to analyse count tables is comprehensively detailed below:

~~~
# **Import .csv files: read counts (count_table.csv), information about samples
and groups (samples_info.csv), and gene annotation file (gene_info.csv)**
> samples_info <– read_csv(“samples_info.csv”)
> count table <– read csv(“count table.csv”)
> gene_info <– read_csv(“gene_info.csv”)
~~~

~~~
# **Call the packages necessary for data processing and analysis**
> library(limma)
> library(edgeR)
> library(genefilter)
> library(pheatmap)
> library(viridis)
> library(dplyr)
~~~

~~~
# **Reformat count_table into a matrix of counts with row.names corresponding
to the column of gene_info which will be used for gene annotation (e.g.
ensembl identifiers). Requires same number of lines in count_table.csv and
gene_info.csv**
> Mat<-as.matrix(count_table[,2:14])
> row.names(Mat)<-count_table$Gene_id
~~~

~~~
# **Create a DGEList object, filtering off the genes not expressed in any dog**
> DGE<-DGEList(counts=Mat, lib.size=colSums(Mat),
norm.factors=rep(1,ncol(Mat)), samples=samples_info, genes=gene_info, remove.zeros=TRUE)
**Removing x rows with all zero counts**
~~~

~~~
# **Transform raw counts into count per million reads (cpm), and log(cpm)**
> DGEcpm<-cpm(DGE, log=FALSE)
> DGElcpm<-cpm(DGE, log=TRUE)
~~~

~~~
# **keep only genes with cpm>1 in at least 3 Dogs**
> keep.exprs<-rowSums(DGEcpm>1)>=3
> DGE<-DGE[keep.exprs, keep.lib.sizes=FALSE]
> DGElcpm<-cpm(DGE,log=TRUE)
~~~

~~~
# **incorporate normalization (scaling) factors to reduce heteroscedasticity,
using the trimmed mean of M-values (TMM) method**
> DGE<-calcNormFactors(DGE, method=“TMM”)
~~~

~~~
# **Genewise Negative Binomial Generalized Linear Fitting using the edgeR package**
> design<-model.matrix(~0+Treatment)
> colnames(design)<-gsub(“Treatment”, ““, colnames(design))
# **‘Treatment’ corresponds to the name of the column in the “samples_info.csv”
file on which the statistical comparison is performed.**
> estdisp<-estimateDisp(BicepsDGE, design)
> fit<-glmFit(estdisp, design)
~~~

~~~
# **Factors=c(“AAVHigh”,”AAVLow”,”AAVMid”,”Healthy”,”None”)**
> lrt<-glmLRT(fit, contrast=c(0,0,0,-1,1)) **# hence contrast=None-Healthy**
~~~

~~~
# **Subset DE genes: p value ≤ 0.05 & fold change ≥ 2 or ≤ -2**
> DE_genes<-lrt[which(lrt$table$PValue<=0.05 & abs(lrt$table$logFC)>1),]
~~~

~~~
# **Same analysis done for contrast=c(1,0,0,0,-1), named lrt2 and DE_genes2
and c(1,0,0,-1,0), named lrt3 and DE_genes3**
~~~

~~~
# **Extract list of Rescued genes in AAVHigh dogs = common genes between
DE_genes and DE_genes2 that are not present in DE_genes3**
> RescHigh<-dplyr::semi_join(DE_genes$genes, DE_genes2$genes, by=“X1”)
> RescHigh<-dplyr::anti_join(RescHigh, DE_genes3$genes, by=“X1”)
~~~

~~~
# **Extract list of Resistant genes in AAVHigh dogs = common genes between
DE_genes1 and DE_genes3 that are not present in DE_genes2**
> ResistHigh<-dplyr::semi_join(DE_genes$genes, DE_genes3$genes, by=“X1”)
> ResistHigh<-dplyr::anti_join(ResistHigh, DE_genes2$genes, by=“X1”)
~~~

~~~
# **Extract list of Induced genes in AAVHigh dogs = common genes between
DE_genes2 and DE_genes3 that are not present in DE_genes**
> IndHigh<-dplyr::semi_join(DE_genes2true$genes, DE_genes3$genes, by=“X1”)
> IndHigh<-dplyr::anti_join(IndHigh, DE_genes$genes, by=“X1”)
~~~

~~~
# **Extract list of DE genes that are present in DE_genes, DE_genes2, and
DE_genes3.**
> TriHigh<-dplyr::intersect(DE_genes$genes, DE_genes2$genes)
> TriHigh<-dplyr::intersect(TriHigh, DE_genes3$genes)
> length(TriHigh$X1)
~~~

~~~
# **Create a matrix with grouped median lcpm values of TriHigh genes**
> TriHighMat<-matrix(c(1:12),nrow=4,ncol=3)
> row.names(TriHighMat)<-TriHigh$X1
> colnames(TriHighMat)<-c(“Healthy”,”XLMTM”,”AAVHigh”)
> for(i in 1:4){
+      TriHighMat[i,1]<-(DGElcpm[TriHigh[i,1],1]+DGElcpm[TriHigh[i,1],7])/2
+      TriHighMat[i,2]<-
(DGElcpm[TriHigh[i,1],4]+DGElcpm[TriHigh[i,1],8]+DGElcpm[TriHigh[i,1],10])/3
+      TriHighMat[i,3]<-(DGElcpm[TriHigh[i,1],12]+DGElcpm[TriHigh[i,1],13])/2
+      }
~~~

~~~
# **Use operators to extract lists of Partially Rescued and Worsened genes**
> PartHigh<-which((TriHighMat[,1]<TriHighMat[,3] & TriHighMat[,3]<TriHighMat[,2]) | (TriHighMat[,1]>TriHighMat[,3] & TriHighMat[,3]>TriHighMat[,2]))
> WorHigh<-which((TriHighMat[,1]<TriHighMat[,2] &
TriHighMat[,2]<TriHighMat[,3]) | (TriHighMat[,1]>TriHighMat[,2] & TriHighMat[,2]>TriHighMat[,3]))
~~~

~~~
# **Similar analyses can be performed for AAVMid and AAVLow dogs by changing
the contrasts taken as inputs for lrt2 and lrt3**
~~~

Gene ontology analyses were performed on the GO consortium online platform (http://geneontology.org/), using the statistical overrepresentation test and the Bonferroni correction for multiple testing. Enrichments with a corrected P value lower than 0.05 were considered significant.

### Data representation

Heatmaps and PCA plots were automatically generated in R-Bioconductor. Other data representations were realized on Graphpad PRISM 7, and Figures were assembled using the vector graphics Inkscape software. The color palette used in this manuscript to distinguish dog groups was extracted from the viridis package on R-Bioconductor and was carefully selected to be easily interpretable by individuals with colorblindness. (https://cran.r-project.org/web/packages/viridis/vignettes/intro-to-viridis.html).

### Data availability

The RNA-Sequencing data from this publication have been deposited to the Sequence Read Archive (SRA) database (https://www.ncbi.nlm.nih.gov/sra), and assigned the Bioproject ID PRJNA49575. The read files are available at the following link: https://trace.ncbi.nlm.nih.gov/Traces/sra/sra.cgi?study=SRP165150

## Acknowledgements

### Funding sources

NIH/NIAMS R01-HL115001, Will Cure Foundation, Joshua Frase Foundation, Association Française contre les Myopathies (AFM-Telethon), Myotubular Trust UK.

### Author contributions

J.-B.D. designed and performed the experiments, analysed the data and wrote the manuscript.

J.G. designed the experiments, analysed the data and assisted in drafting the manuscript.

M.W.L. processed the canine muscle samples and assisted in drafting the manuscript.

R.W.G. performed the assessments of muscle force and assisted in drafting the manuscript.

J.T.G. designed the experiments and assisted in drafting the manuscript.

A.B.-B. provided the gene therapy vector, designed the experiments and assisted in drafting the manuscript.

M.K.C. designed the experiments

D.L.M. designed the experiments, analysed the data and wrote the manuscript.

### Conflict of interest

MWL is a member of advisory boards for Audentes Therapeutics, Solid Biosciences, and Ichorion Therapeutics. He is also a consultant for Wave Life Sciences and Dynacure.

A.B.-B. and M.K.C. are inventors of a patent on gene therapy for myotubular myopathy.

### The Paper Explained

#### PROBLEM

Gene therapy using recombinant adeno-associated virus (rAAV) vectors is an attractive approach to treat neuromuscular disorders resulting from single gene mutations. Despite encouraging results in large animal models and in recent clinical trials, gene therapy is sometimes considered a “hit-or-miss” technology, because the exact molecular mechanisms driving disease rescue remain elusive, and biomarkers of rAAV corrective impact in target cells are missing.

#### RESULTS

In this study, the complete rescue of X-linked myotubular myopathy (XLMTM) after rAAV gene therapy in dogs is used as a model to develop analytical tools and help decipher the impact of rAAV on the transcriptome. We first proved that transcriptome analysis by RNA-sequencing successfully discriminates dogs based on the extent of disease correction. It confirms previously described gene deregulations associated with XLMTM phenotypes in murine models and points towards new players in disease development. From these results, we created an analytical framework based on a defined terminology, statistical comparisons, and intuitive visualization tools to precisely characterize the remodelling of the transcriptome induced by gene therapy. In line with the recovery of muscle structure and function highlighted in our previous study, the indicators included in this pipeline prove that XLMTM correction also occurs at the level of the transcriptome in a dose-dependent manner.

#### IMPACT

Our methodology tackles an important question in the field of gene therapy, related to the molecular mechanisms of disease reversal. It provides a way to identify clusters of genes or molecular pathways positively affected by the treatment, and to highlight those escaping correction as potential targets for combination treatments. It can also predict the identity of interesting biomarkers to test in future preclinical and clinical settings. Given the increasing influence and decreasing cost of RNA-sequencing technologies, we anticipate that this type of analysis will become part of the standard procedures as gene therapy is rapidly moving to the clinic.

