## Supplementary material for "AAV-mediated gene transfer restores a normal muscle transcriptome in a canine model of X-linked myotubular myopathy"

**Supplementary Figure 1: Gene ontology analysis of the TOP 500 most informative genes.**

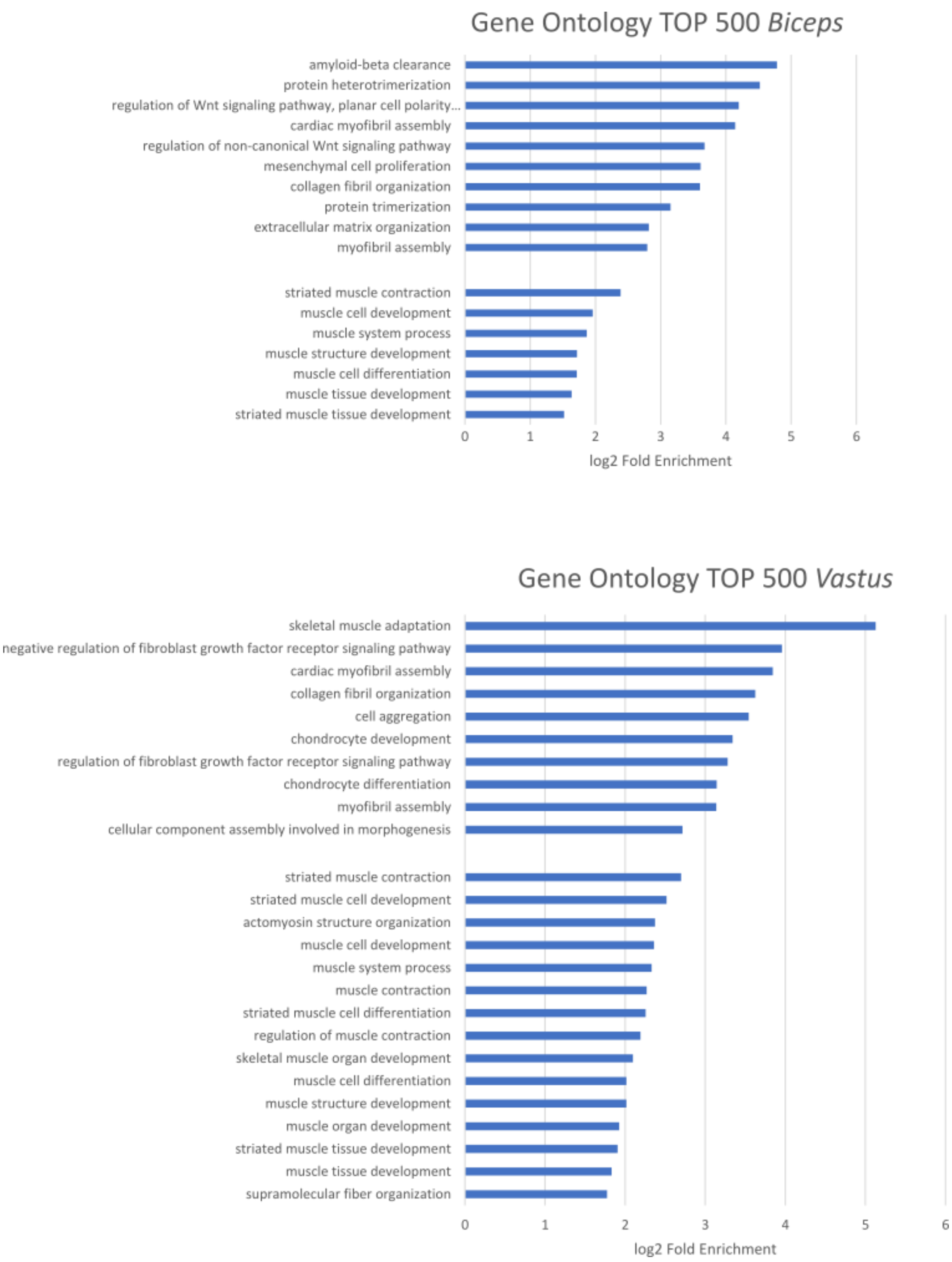


Gene ontology analysis of the 500 genes with the highest expression variance across samples in the *Biceps femoris* (Cummings et al.) and *Vastus lateralis* (bottom). The 10 GO terms with the highest fold enrichment (FE) are represented, and the muscle-related terms with a log2 FE > 1.

**Supplemenatary Figure 2: Over-expression of developmental genes in XLMTM muscles.**

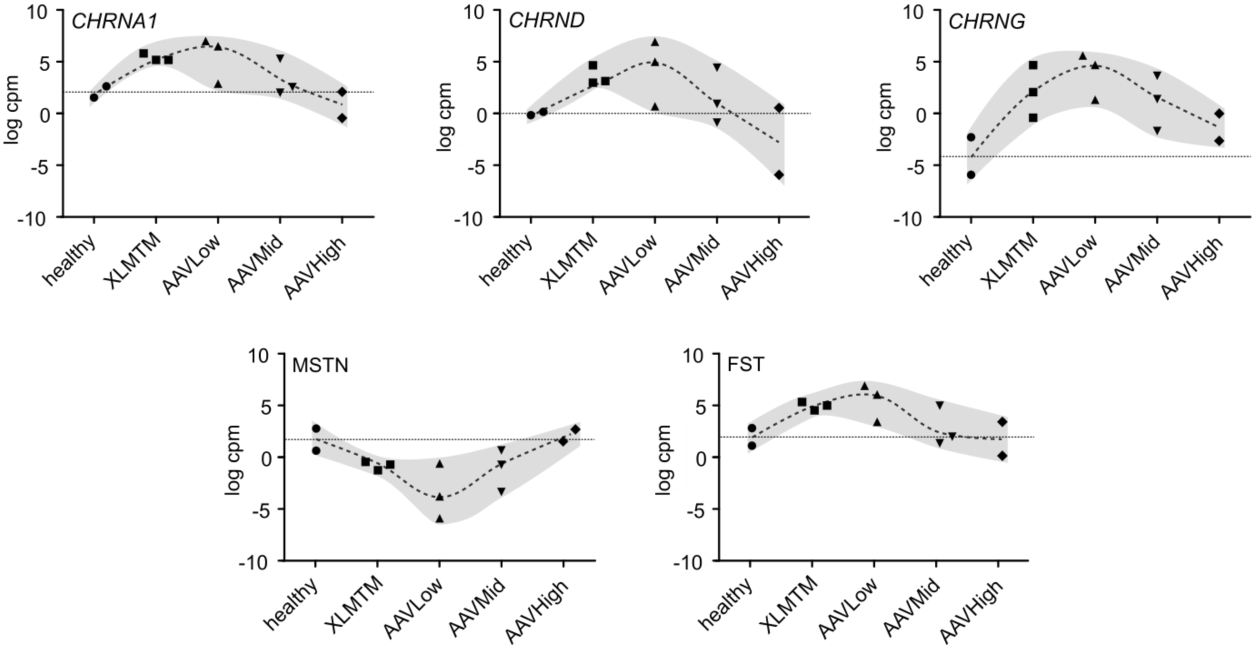


Gene expression data (log cpm) for three genes encoding cholinergic receptor subunits (*CHRNA1*, *CHRNG*, *CHRND*) and two myokines: myostatin (*MSTN*) and follistatin (*FST*) in the *Biceps femoris*. Each symbol represents one individual dog. The median expression in healthy controls as a horizontal dashed line, the expression range as a shaded area, and the evolution of the median as a dashed curve. cpm: count per million reads, log: base 2 logarithm.

**Appendix Figure 3: Relative expression of the common candidate biomarker genes.**

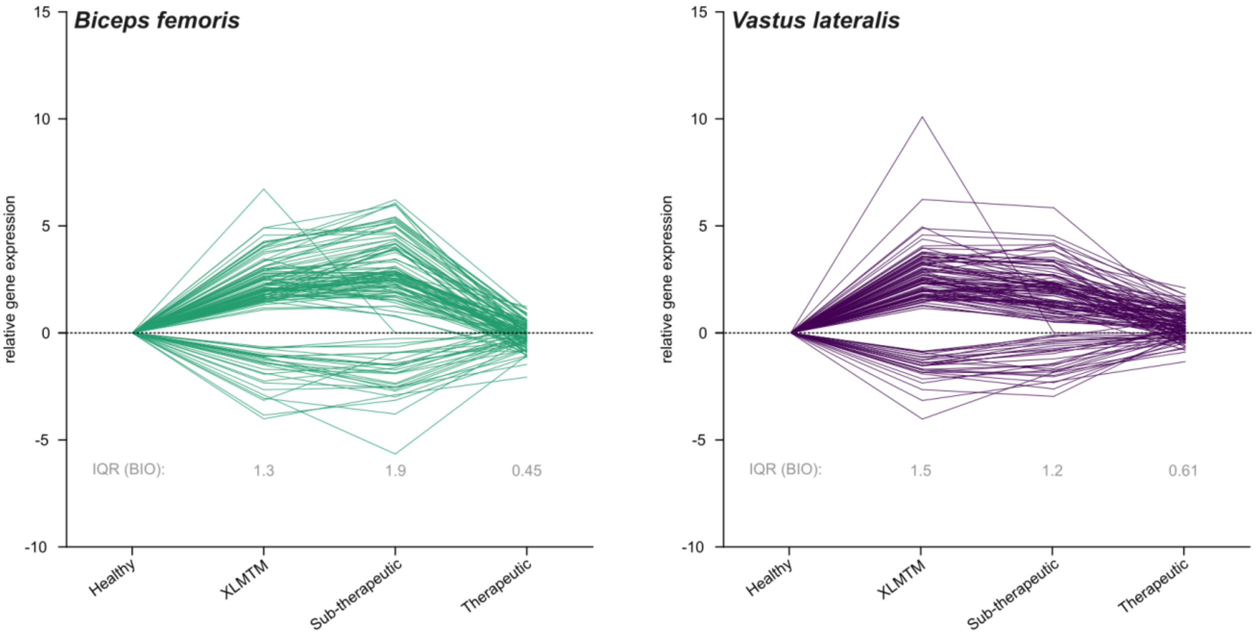


Relative expression of the 120 candidate biomarker genes expressed in log fold change (log FC) after normalization with healthy controls. The interquartile ranges were determined from positive log FC values (negative log FC were converted into absolute values). The dashed line at Y = 0 represents no variation in gene expression from healthy controls.
